# Simple post-translational circadian clock models from selective sequestration

**DOI:** 10.1101/2020.02.21.958827

**Authors:** Mark Byrne

## Abstract

It is possible that there are post-translational circadian oscillators that continue functioning in the absence of negative feedback transcriptional repression in many cell types from diverse organisms. Apart from the KaiABC system from cyanobacteria, the minimal molecular components and interactions required to potentially create “test-tube” circadian oscillations in different cell types are currently unknown. Inspired by the KaiABC system, I provide proof-of-principle mathematical models that a protein with 2 (or more) modification sites which selectively sequesters an effector/cofactor molecule can function as a circadian time-keeper. The 2-site mechanism can be implemented using two relatively simple coupled non-linear ODEs in terms of site occupancy; the models do not require overly special fine-tuning of parameters for generating stable limit cycle oscillations.

## I. INTRODUCTION

There are circadian (≈ 24 hr) clocks in most organisms [1]. These internal biological clocks are experimentally characterized by sustained oscillations in one or more measured outputs under (approximately) constant conditions (e.g., constant light or darkness). Circadian oscillators can also be entrained via external stimuli, such that the internal oscillator’s period and relative phase can be synchronized by the entraining stimulus [2]. In contrast to general biochemical oscillators, the oscillatory period of a circadian clock remains almost unchanged for different constant temperatures in the relevant physiological range (temperature compensation) [3]. The simplest known example of a circadian clock that functions outside cells (in a “test-tube”) is the 3-protein KaiABC system [4, 5]. This type of clock is termed a post-translational oscillator (PTO) since its operation does not require dynamical feedback from a genetic circuit. In contrast, the fundamental oscillatory design of the circadian clock in most organisms in vivo is believed to consist of a transcription-translation negative feedback loop (TTFL) which is influenced by post-translational modifications (see, for example [6–8]). However, negative feedback TTFLs do not generally imply sustained oscillations [9, 10]. In this context, mathematical models were originally required as proof-of-principle that the TTFL mechanism was quantitatively viable, in the sense that simple TTFL mathematical models could, in principle, reproduce experimentally realistic circadian oscillations in the relevant measured outputs [11].

Both simplified (2D) and more complex mathematical models of eukaryotic TTFL circadian clocks have been investigated [12, 13]. However, there is evidence of the existence of circadian PTOs in a variety of species absent genetic feedback loops [14–16]. These findings suggest that it may be useful to investigate post-translational designs which can satisfy the defining conditions for a circadian clock [17]. The elucidation of basic molecular designs which can create circadian clocks can presumably serve as a guide to experimental searches for possibly other protein-based PTOs and for investigating the apparent degree of simplicity (or difficulty) required for intracellular post-translational molecular methods of timing.

In particular, finding simplified two dimensional mathematical models of circadian clocks is potentially useful since a variety of mathematical tools (phase plane analysis, Hopf bifurcation theory) can be used to analyze and visualize the system’s dynamics. This study demonstrates the existence of two simple designs whose mathematical models are perhaps the simplest 2D circadian clock models for post-translational molecular oscillators. Both designs require a protein with at least two regulatory sites, selective sequestration of an effector molecule by one of the protein states, and a separation into fast-slow kinetics for regulation of site occupancy. These general conditions are sufficient to generate stable limit cycle oscillations in the population occupancy of the sites for reasonable rates and parameter values. Furthermore this class of “selective sequestration” models permits both entrainment and temperature compensation, by assuming one of the sites (the “fast” site) is sensitive to external perturbation while the “slow” regulatory dynamics on the other site is essentially buffered from external perturbations.

## II. RESULTS

### 1. Post-translational oscillatory mechanisms from two protein sites and selective sequestration

For simplicity assume a “core” clock protein (*X*) with two modification sites (*a* = 1 and *b* = 2) in which the rate of modification of the residues can be altered by another “effector” molecule (*Y*). For the sake of generality we suppose the molecule that is transferred to each site of the protein remains unspecified (e.g., a phosphoryl group, ATP, or an oxygen atom) - for the KaiC protein there are two phosphorylation sites per KaiC monomer and the rate of (auto) phosphorylation of the sites is affected by another protein (KaiA) in a hyperbolic manner [18, 19]. We propose two general simple designs for an autonomous clock using sequestration of *Y* based on the occupancy of the two sites (Fig 1).

**FIG. 1.**
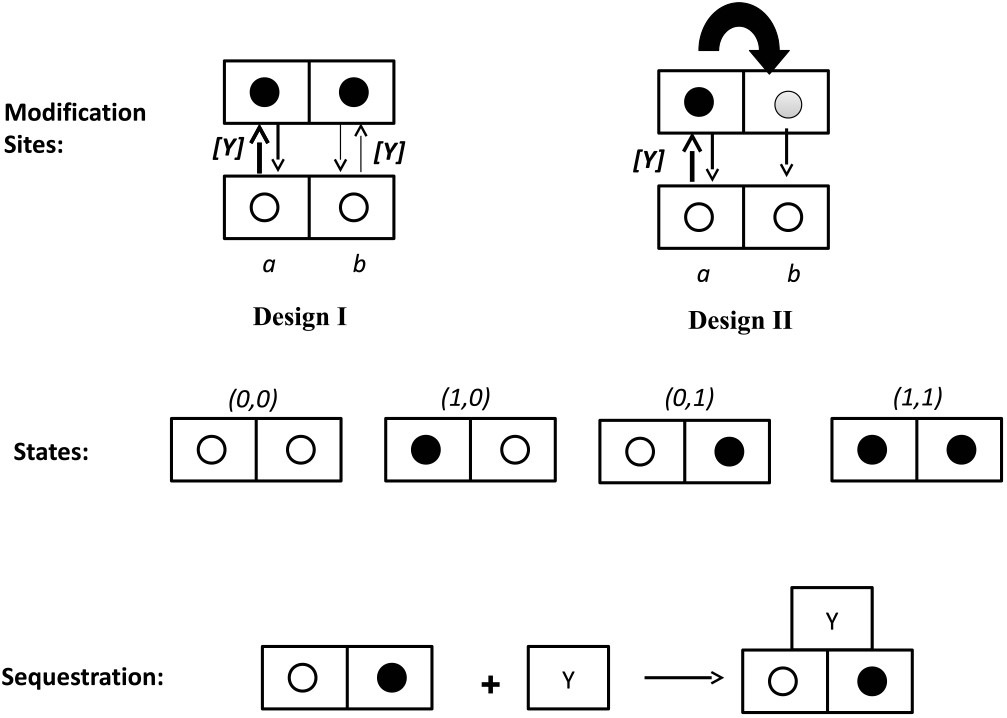
Two possible simple designs for a protein-based clock. In design I, two modification sites on the protein *X* (not necessarily adjacent) are both regulated by the effector, *Y*. Filled circles indicate occupancy of a protein site, by addition of a small molecule to that residue (for example). In design II the occupancy of one site (regulated by *Y*) is transferred to a second site. Only one of the four protein states sequesters the effector molecule, Y.)

In one oscillatory scheme each site is independently modified and the kinetics proceed at different rates on the two sites; in a second class of models the occupancy of one site is transferred to the other site via some unspecified mechanism (e.g., an intra-protein transfer). Assume the population kinetics of the two sites follows 1st order kinetics so that the fractional occupancy of each site (0 < *x_i_* < 1, e.g., degree of “phosphorylation”) in the protein population is described as follows:

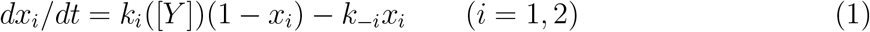

where *i* labels each modification site and a constant time-independent “decay” (e.g., “dephosphorylation”) term is allowed in the kinetics (a derivation of Eq. (1) in terms of protein abundance and site occupancy is given in the supplemental information). In general the modification rate(s) on the site(s) are assumed to vary as smooth, monotonic functions, *f*, of the effector concentration, which are parameterized using *k_i_* = *k_i,max_f* ([*Y*]) where 0 < *f* ([*Y*]) < 1; hyperbolic and linear regulatory functions of the rates are examined below.

An individual protein (*X*) with two modification sites can be in one of four states: (*a, b*) = (0, 0), (1, 0), (0, 1), (1, 1) with zero indicating an un-occupied site and one occupied. Population statistical arguments (see supplemental) in a well-mixed population imply the average fraction of proteins in each of the four states at time *t* is

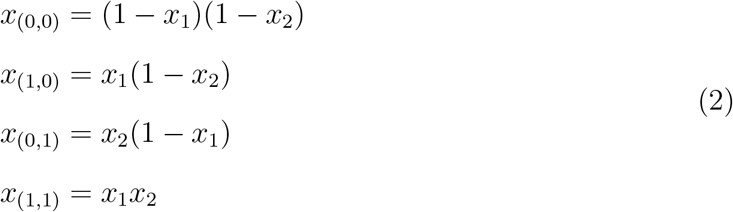

Thus the relative protein abundances of the four states can be computed after solving Eqn 1. Now suppose the modification rates (*k_i_*) of both sites follow Michealis-Menten regulatory kinetics, depending hyperbolically on the concentration of the effector molecule (*Y*):

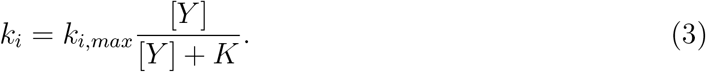

Letting site *a* represent the ‘fast’ site (*k*_1*,max*_ >> *k*_2*,max*_) breaks the symmetry in the model and sequestration of *Y* by the (0, 1) protein states can generate sustained oscillations depending on the rate constants and model parameters. A simple “rigid” model of sequestration is to instantaneously alter the concentration of *Y* according to the concentration of *X* proteins in one of the four protein states [19]. If an average of *n* molecules of *Y* are sequestered per *X* protein in the state (0, 1), the non-sequestered concentration of [*Y*] a time *t* is given by:

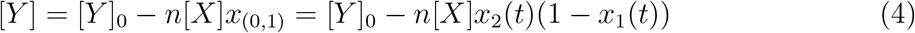

with the physical constraint that [*Y*] ≥ 0; [*Y*]_0_ indicates the initial concentration of *Y* (a more complete derivation is given in supplemental information). Equivalently we are assuming the rate of change of free effector (*Y*) due to sequestration (or “inactivation”) of *Y* is directly proportional to the (negative) rate at which the (0, 1) protein states are produced or degraded so the relevant timescale for *Y* sequestration/inactivation is assumed to be much shorter than the circadian timescale.

For this mechanism, the dependence of the system dynamics in terms of the model parameters and rates is best seen by rescaling equations (2) and (4) by the fixed protein concentration [*X*] and the parameter *n*:

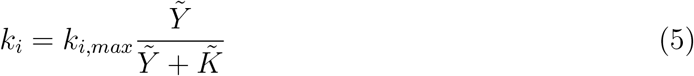

and

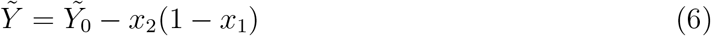

where 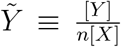 is the dimensionless scaled fraction of the effector, *Y* (non-negative); 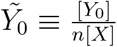 is the equivalent dimensionless scaled fraction of initial effector [*Y*_0_], and 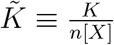 is a dimensionless Michaelis constant. The system dynamics is encoded by the two fixed parameters (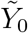 and 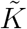) and three (ratios) of rate constants (since one rate constant can be absorbed into a dimensionless time parameter by re-scaling both differential equations in Eqn.(1) by, e.g., 1*/k*_1*,max*_). For example, sustained ≈ 24 hr oscillations are reproduced using the parameters 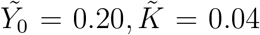 and the rates *k*_1*,max*_ = 0.4*hr*^−1^ = *k*_−1_, *k*_2*,max*_ = 0.03*hr*^−1^ = *k*_−2_ (Fig 2). Numerical integration suggests a limit cycle upon varying the initial conditions (Fig 3A). Sustained oscillations are possible as a result of an autocatalytic negative feedback loop (from sequestration), previously suggested in several mathematical models of the KaiABC clock ([20],[19],[21]). In this 2D model the qualitative description for sustained oscillations is that the slow kinetics on one site allows the site with rapid kinetics to reach near maximal occupancy before sequestration takes effect. Once sequestration starts to occur from the slow formation of (0, 1) states, loss of occupancy on the fast site drives the transition (1, 1) → (0, 1). This causes further sequestration of *Y* and increases the rate of further loss of occupancy on the fast site creating additional (0, 1) states (auto-catalysis). Slow loss of occupancy from the 2nd site assures that the 1st site becomes largely unoccupied until de-sequestration of *Y* occurs simultaneous with the transition (0, 1) → (0, 0). Numerical investigations of the solution space of this model (Eqs. 1, 5 and 6) indicate that the existence of oscillations is quite sensitive to the parameters 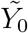 and 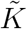 (Fig 3D). Qualitatively, as the relative concentration of initial effector is lowered (for constant 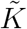) the oscillatory period decreases because sequestration of *Y* takes less time; for fixed initial effector, increasing the Michaelis constant delays the onset of sequestration resulting in a longer oscillatory period. Numerical solutions indicate that sustained oscillations require a “fast-slow” separation of timescales for the maximum modification rates on the two sites (Fig 3B). Varying both *k*_1*,max*_ and *k*_2*,max*_ over the range [0, 1] *hr*^−1^ in steps of 0.01 *hr*^−1^ (and setting *k*_−1_ = *k*_1*,max*_, *k*_−2_ = *k*_2*,max*_) suggests sustained oscillations occur when *k*_2*,max*_ < 0.1*k*_1*,max*_ (Fig 3B). As expected the oscillatory period is only weakly dependent on *k*_1*,max*_ but strongly dependent (power-law dependent with a power ≈ −0.75) on *k*_2*,max*_ (Fig 3C), which is important in interpreting both temperature compensation and entrainment in these clock models.

**FIG. 2.**
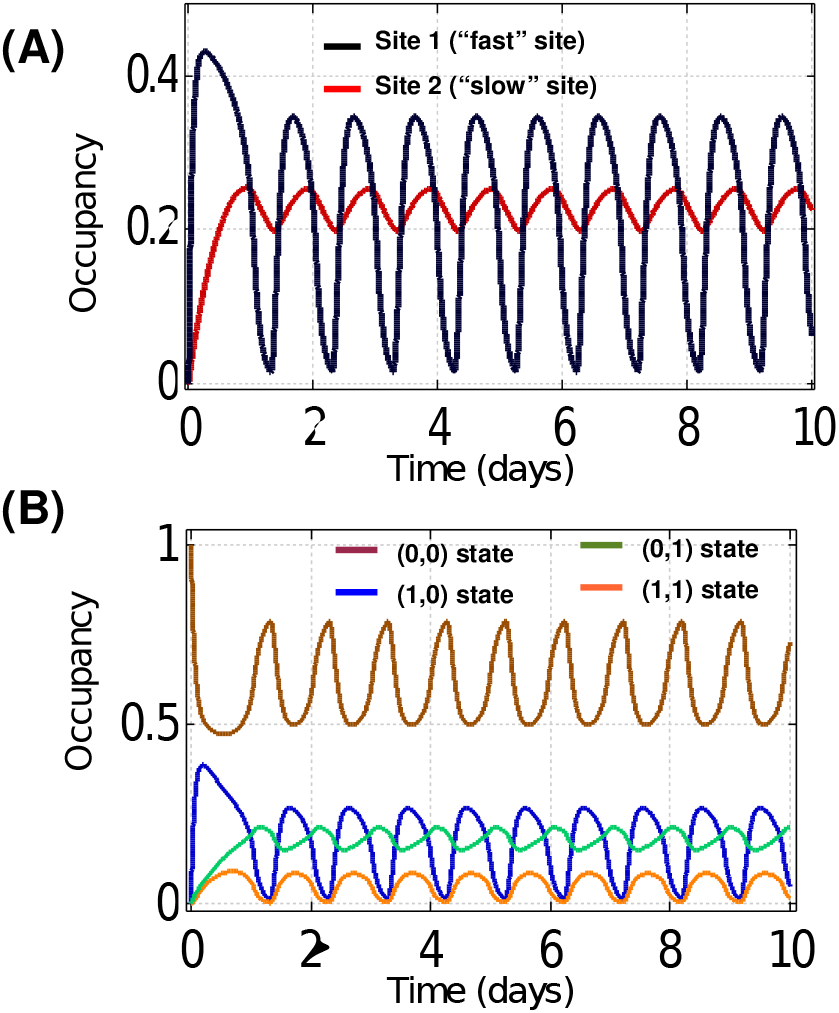
Design I with independent (hyperbolic) modification of the sites by *Y*. (A) Example population occupancy kinetics for the two protein sites (rates and parameters are in the main text). (B) Dynamics for the corresponding 4 protein states.

**FIG. 3.**
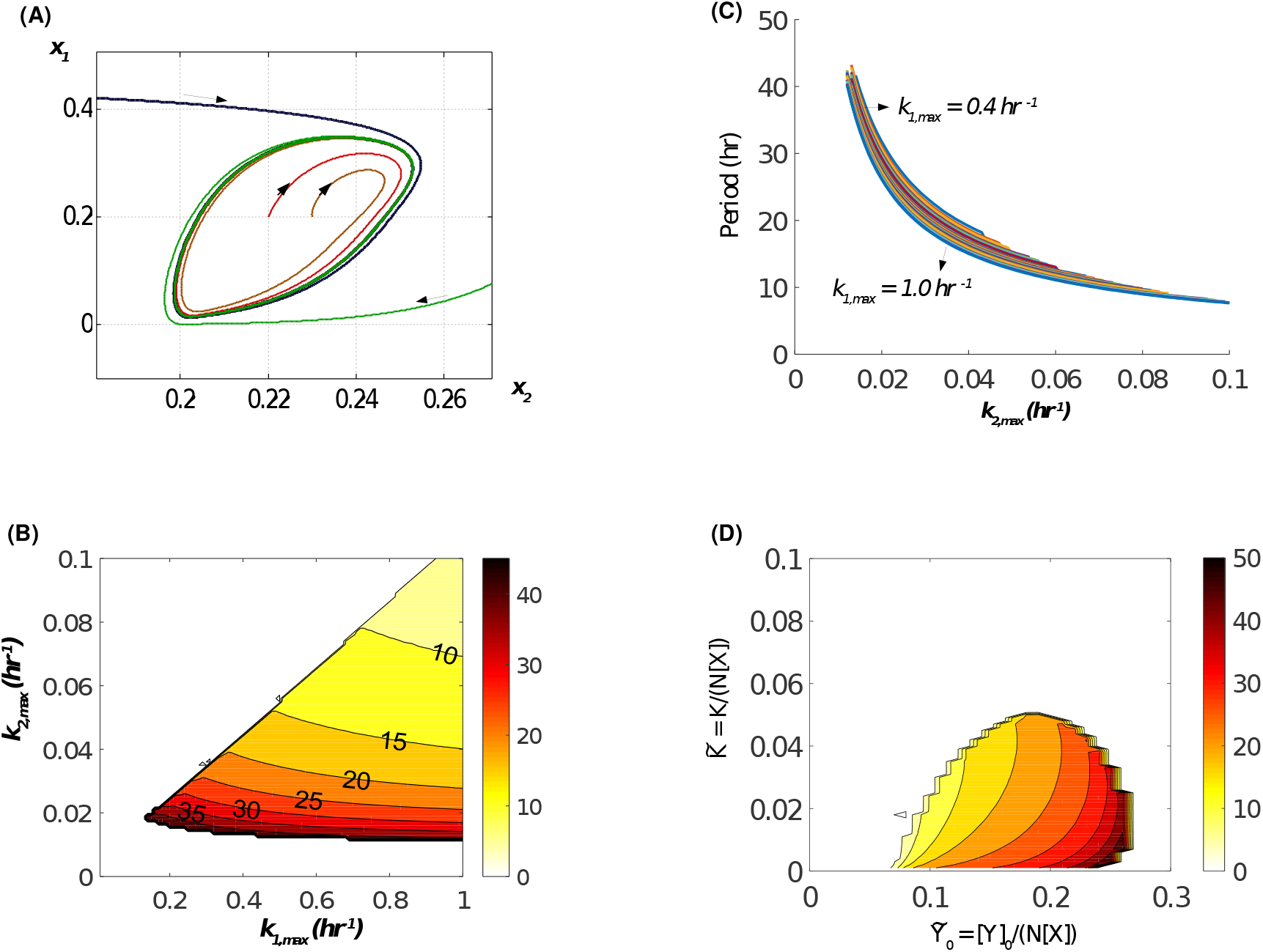
Model I: (A) Stable sample limit cycle trajectories for different initial conditions using the rates and parameters from fig 2. (B) Parameter region allowing oscillations and the oscillatory period in hours assuming 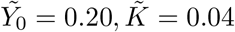 (C) An elaboration of Panel (B), indicating sensitive dependence of period (power-law) to the “slow” rate constant for site 2 modification (D) Parameter region allowing oscillations and the oscillatory period in hours for a given dimensionless Michaelis constant and scaled initial concentration of effector, assuming *k*_1*,max*_ = 0.4*hr*^−1^ = *k*_−1_, *k*_+2*,max*_ = 0.03*hr*^−1^ = *k*_−2_.

Interestingly there is an even simpler class of sequestration models that can generate stable limit cycles (and might be employed biochemically). Consider the “transfer” design in which the molecule occupying site 1 is transferred to site 2 at rate *k_T_* so that the model is now described by a class of ODEs:

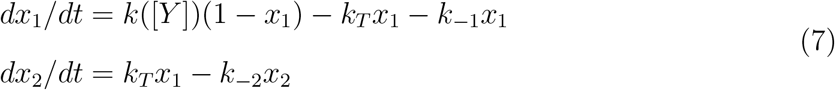

where terms for both transfer and loss of occupancy for site 1 have been included. In these designs the rate *k* can depend linearly on the substrate 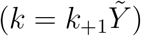 and generate limit cycle oscillations (hyperbolic variation can also generate stable limit cycles). Non-dimensionalizing (*τ* ≡ *k*_+1_*t*) and neglecting loss of occupancy without transfer (for the moment), *k*_−1_ = 0, gives the following simple system 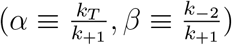, where:

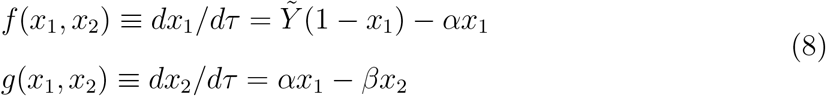

with 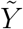 given by Eqn.(6), 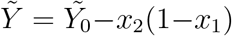 and 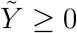. For example there are stable limit cycle oscillations for *α* = 0.25*, β* = 0.1, 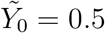; setting *k*_+1_ = 0.7*hr*^−1^ generates a circadian timescale for the oscillations (Fig 4, 5A). Allowing loss of occupancy from site 1 lowers the oscillation amplitude and can result in damped oscillations; examples for *k*_−1_ = 0.01*hr*^−1^ and *k*_−1_ = 0.05*hr*^−1^ are shown in Fig 5B.

**FIG. 4.**
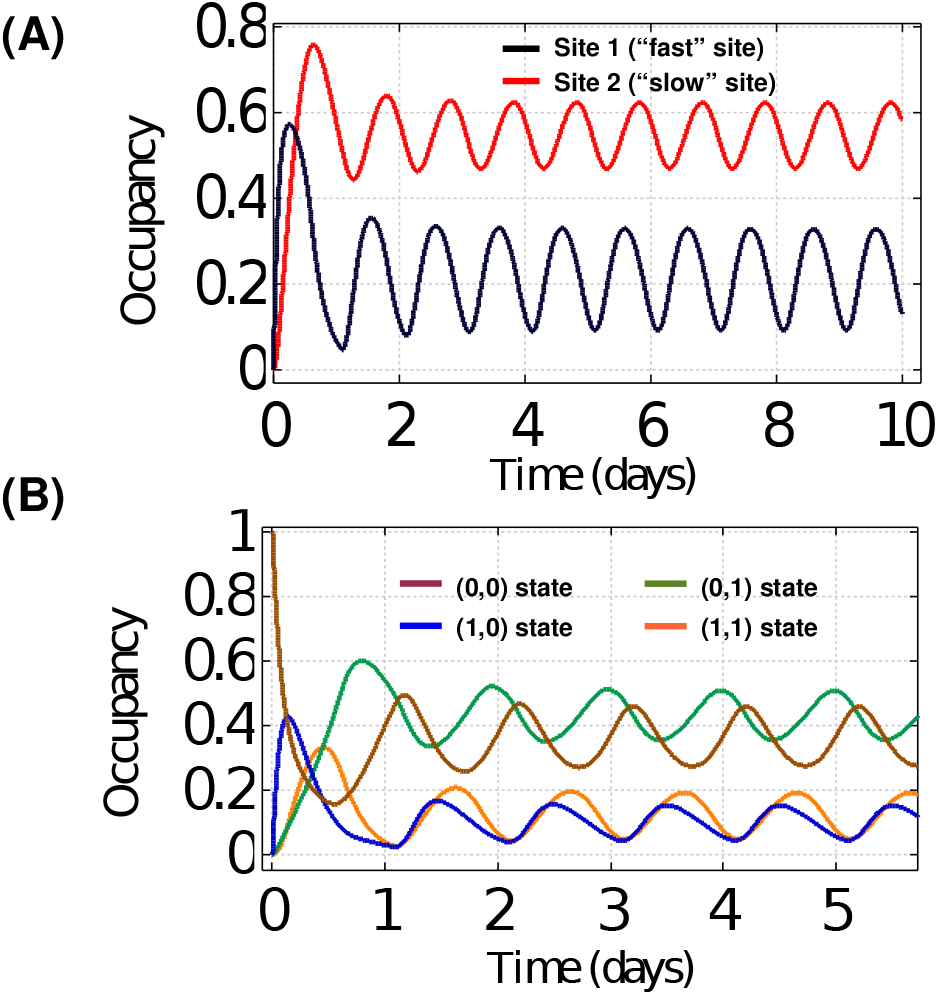
Design II with transfer of occupancy from site 1 to site 2. (A) Example population occupancy dynamics for the two protein sites (rates and parameters are in the main text). (B) dynamics for the corresponding four protein states.

**FIG. 5.**
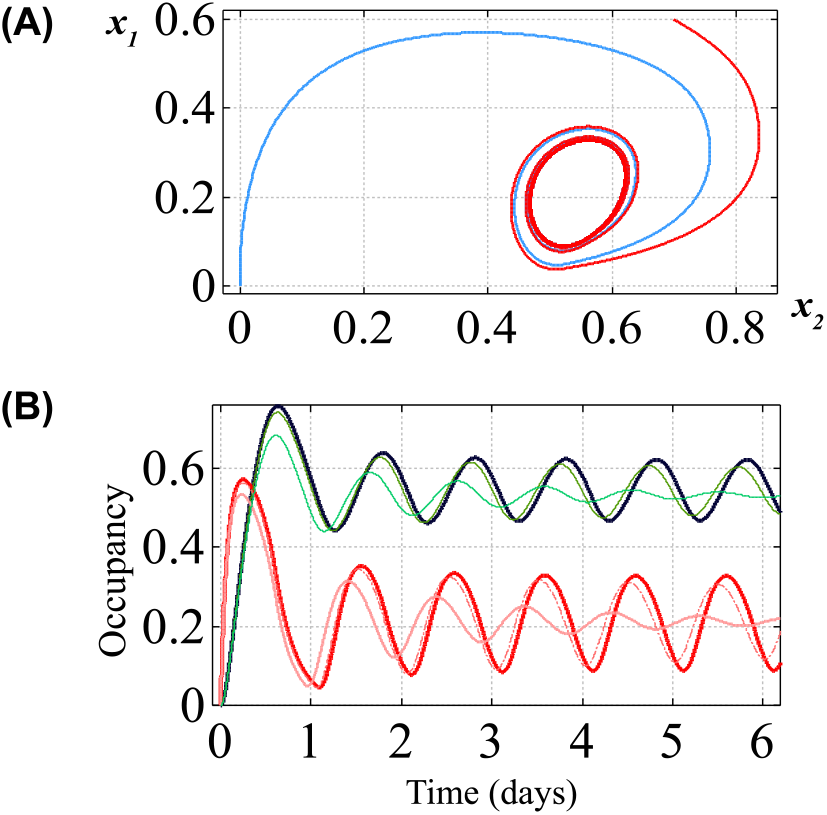
Design II: (A) Stable sample limit cycle trajectories for different initial conditions using the rates and parameters from fig 4. (B) Allowing both loss of occupancy from site 1 and transfer to site 2; *k*_−1_ = 0.05*hr*^−1^ (thin trace, damped oscillations) and *k*_−1_ = 0.01*hr*^−1^ with slightly reduced amplitudes compared to Fig 4A traces (thicker red and black)

The simple dynamical system can be studied analytically - see Supplemental for a linear perturbative analysis about the steady-states. Direct integration of the ODEs confirms the regions of instability and oscillatory period estimates from a linear stability analysis of the dynamical system (Fig 6). As an example, consider 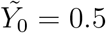. Linear (in)stability constraints suggest oscillations generally require *β* < 0.1*, α* < 0.5 and *β < α* (Fig 6). The dimensionless oscillatory period 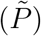 is fit well (*R*^2^ > 0.99) by different power-law functions for a given *β*:
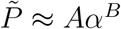 with (*A, B*) = (2.74, −1.53) for *β* = 0.01; (*A, B*) = (3.37, −1.23) for *β* = 0.05; and (*A, B*) = (2.89, −1.27) for *β* = 0.1. The physical period is 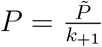 which, in this regime of parameters for oscillations gives an approximate physical oscillatory period:

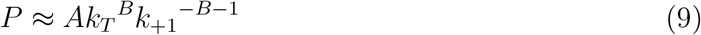

**FIG. 6.**
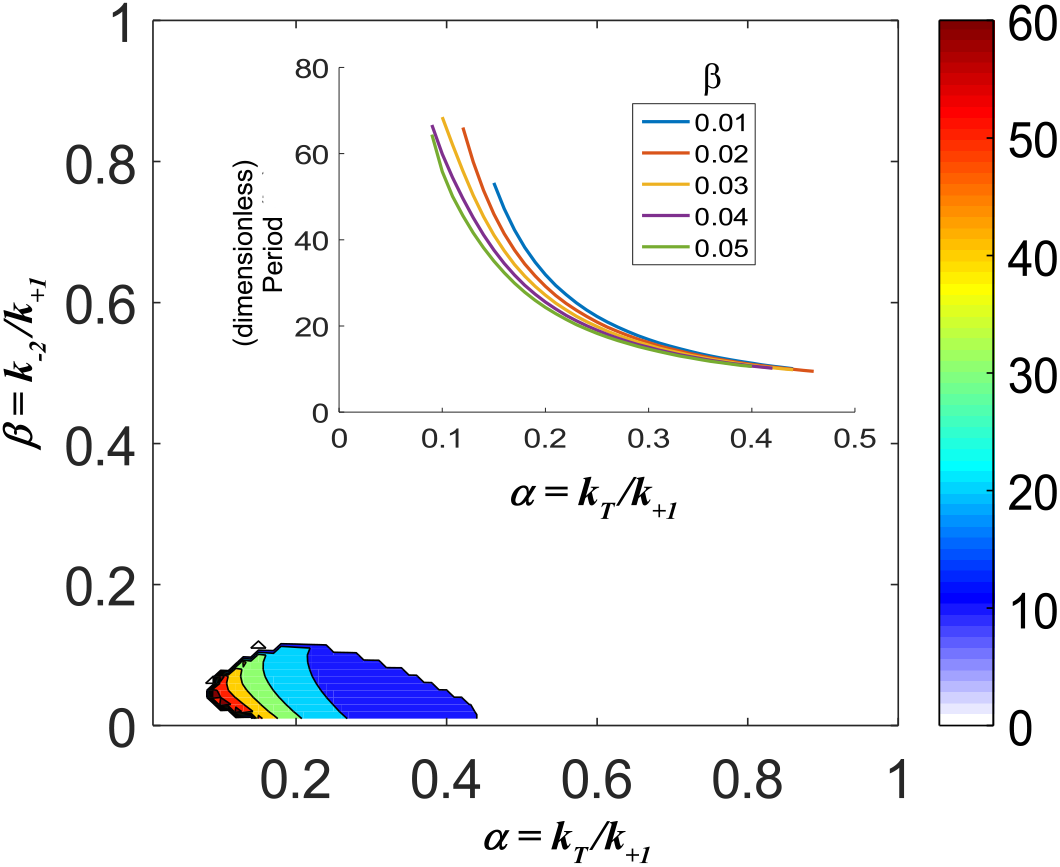
Approximate region of stable limit cycle oscillations for design II from numerical integration; colormap shows dimensionless period as the ratio of rates are varied (fixed 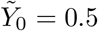) and the inset shows the approximate power-law dependence of the dimensionless period on *α*.

For example, for *β* = 0.1:

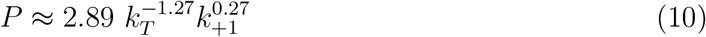

Thus the oscillatory period in this class of models is primarily set by the “transfer” rate (*k_T_*).

### 2. Entrainment and Temperature Compensation

Circadian clocks are both sensitive to external perturbations (can be entrained) yet retain near period invariance under varied environmental conditions, such as temperature fluctuations. In these simple models we can suggest a possible mechanism; the oscillator period is “mildly” sensitive to external perturbations so that the oscillation period is approximately constant and set by the “slow” internal dynamics, while the “fast” dynamics (e.g., *k*_+1_) incorporates external perturbations. In this model oscillator, all perturbations (metabolic, light or dark pulses, temperature, etc.) interact only through the modification rates, assuming the perturbation does not alter the relative protein abundances 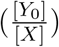. In these models the change in period (*δP*) in terms of any perturbation in rates (*δk_j_*) is, to 1st order,

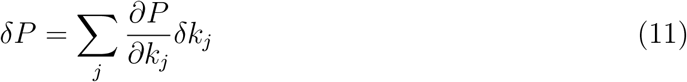

This perturbation should be approximately zero for “compensation” to be effective (temperature or other perturbations). There are two generic possibilities; one is that the effective rates in the model are “trivially” insensitive to the perturbation (*δk_j_* ≈ 0) due to structural properties of the protein(s). However if the system were completely “trivially” compensated then entrainment would not be possible (except by protein and/or effector abundance variation). The other is that the system is “tuned” to some degree so that one perturbation which tends to increase the period is countered by another that decreases the period ([22]). More generally, in the parameter space of *P* (*k_j_*) approximately flat regions correspond to approximate period-invariant sub-spaces of the parameter space. The approximate power-law dependence (Eqn 9) in design II displays this compensatory mechanism with the constraint that the dynamical system’s period remains invariant when setting the sum of partials to zero in the power-law approximation:

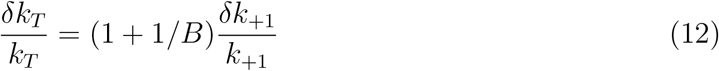

For example, consider initial unperturbed rates of *k*_+1_ = 0.7*hr*^−1^ and *_T_* = 0.2*hr*^−1^ for *β* = 0.04. Numerically the period is *P* ≈ 21.77*hrs*. Simulations suggest a doubling of the site 1 rate (typical for a 10 °C increase), *k*_+1_ = 1.4*hr*^−1^ is almost compensated (*P_new_* ≈ 21.86*hrs*) by about a 30 percent increase in the transfer rate (corresponding to *B* ≈ −1.4, *k_T_* = 0.26*hr*^−1^), Fig 7A. It is clear from direct integration of the model that period compensation is possible as predicted by Eqn 11 (Fig 7A,B); as the fast-rate is further lowered below *k*_+1_ ≈ 0.5*hr*^−1^, the oscillations start damping with increasing period. If we assume the rate doubling on the 1st site corresponds to a 10 °C increase, these parameters give a Q10 (for the period) of about 1.004. Since these are tuned parameters, a desired Q10 can be selected by a judicious choice of rate compensation, corresponding to a choice of activation energy threshold(s) for regulation of site occupancy using the Arrhenius-Boltzmann temperature-dependence of the rates (*k_i_* ∝ *A_i_* exp(−*E_i_/*(*kT*)), as previously described for general biochemical kinetics ([22]).

**FIG. 7.**
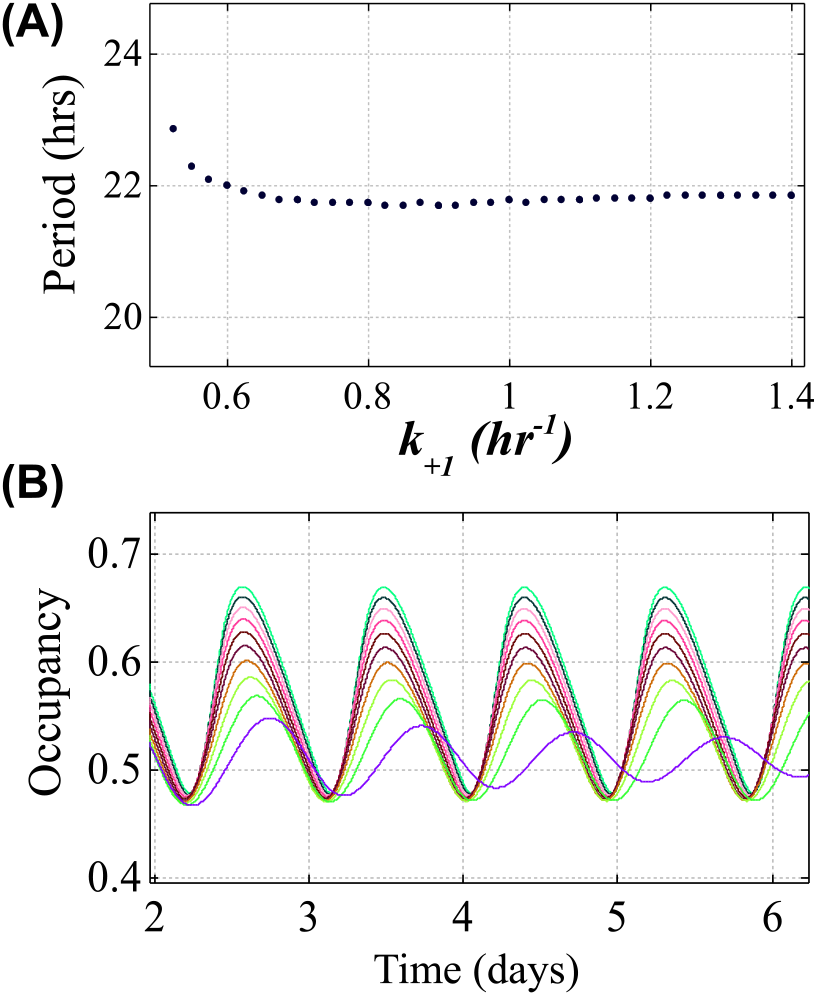
(A). Approximate period invariance of the oscillations as the fast rate (*k*_+1_) is varied while the transfer rate (*k_T_*) is slightly adjusted according to Eqn (12) in the text with *B* ≈ −1.4. (B). Sample site 2 occupancy oscillations corresponding to the variation of *k*_+1_ shown in Panel A (larger oscillation amplitudes correspond to larger *k*_+1_).

Entrainment in the model design II is examined using both continuous and discrete (pulse) perturbations. For small amplitude perturbations, “stable” entrainment is possible within an approximate range of 19 to 26hrs for these parameters. The following external driving function was assumed: 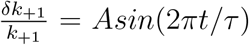 with amplitude, *A* = 0.25 chosen. The transfer rate was modified to retain near period invariance in the absence of the external continuous sinusoidal perturbation (*B* ≈ −1.4 in Eqn 12). For example, the period shifts from 21.77*hrs* (unperturbed) to 24.02*hrs* for an external *τ* = 24 hr rhythm and from 21.77 hrs (unperturbed) to 19.04 hrs for an external *τ* = 19 hr rhythm (Fig 8A). Entrainment is also examined using 2-hr pulse reductions in the rates. Phase response curves (PRCs) are constructed using 2-hr pulses by transiently reducing the the on-rate starting on day 4.5 of the unperturbed oscillator (Fig 8 B,C). The sample PRCs in Fig 8 show reductions of *k*_+1_ = 1.0*hr*^−1^ to {0.0, 0.5, 0.8} *hr*^−1^ starting on day 4.5 of the unperturbed oscillator over one oscillations cycle and computing the phase shift relative to the unperturbed control on day 10 of the oscillation. The transfer rate was also adjusted as previously described according to Eqn 12 using *B* = −1.4. The long “dead phase” of the PRC in these models is because sequestration already abrogates the fast rate (*k*_+1_ ≈ 0) so that further reductions in the rate by an externally applied down-pulse has little effect during the time interval of significant sequestration.

**FIG. 8.**
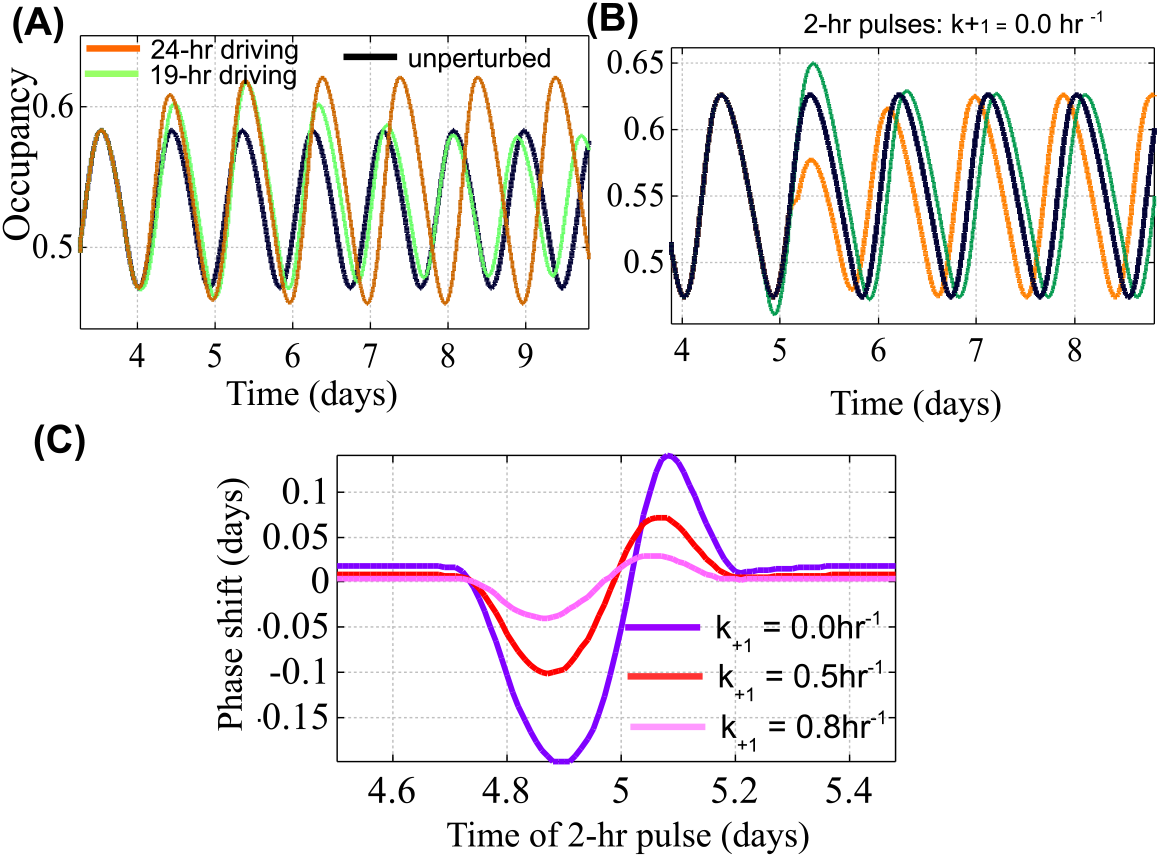
(A). Entrainment to externally driven oscillations of the rate *k*_+1_ starting on day 4; black = control, orange = 24hr, green = 19hr. (B). Sample traces including pulse reductions in the fast rate (setting *k*_+1_ = 0 for 2-hr intervals starting on day 4.5). (C). Sample phase response curves (PRCs) for varying amplitude pulse reductions (measured on day 10) with *k_T_* varied according to Eqn 12.

## III. DISCUSSION

The model designs and simulations in this paper suggest that the molecular interactions required for a post-translational circadian clock could be surprisingly simple. In these designs, selective sequestration of an effector (regulator) of protein site occupancy and a separation of time-scales in the dynamics of regulation of the protein sites appear to be sufficient for generating sustained oscillations. The model can be easily generalized to a protein with *N* regulatory sites of which a subset of the 2^*N*^ states selectively sequesters an effector (or multiple effectors) for finer clock regulation, coupling to other proteins, improved “buffering” of the slow dynamics, etc. Already, with just 2 protein sites and 4 states it is possible to both entrain the oscillations to external cues and have the limit cycle oscillations compensate for fluctuations in the fast dynamical variable (e.g., from temperature or metabolic perturbations). A limitation of the models is that much complexity is encoded in the effective rate constants, including energy regulation and ATP-ADP dynamics which were not included in these models to keep both the state space and parameter space as small as possible. In particular for circadian clocks, the effective rates are much slower than typical rates from enzyme or binding kinetics (*s*^−1^ or less), whereas TTFL models typically incorporate minute to *hr* timescales for significant protein abundance changes. In the Ka-iABC clock, the slow ATPase dynamics of KaiC are implicated in the slow characteristic oscillatory timescale ([23]) and a similar slow ATP hydrolysis would likely be involved in setting the effective rates in these models. Another limitation in these models, also used to reduce the potential state-space of dynamical variables, is that sequestration was not explicitly modeled using mass action (for example); the linear sequestration model by one of the protein states (eqn 6) is an effective and direct method to simulate selective sequestration ([19]) but introduces a hard cutoff in these models reflected in the “fast” rate abruptly shifting to zero.

A prediction of the simplest 2-site/4-state version of the model is that evolutionary mechanisms should have selected rates such that the existence of oscillatory behavior is robust under typical fluctuations of the effective rates; the “transfer” model predicts the most likely values of *α* in the range 0.1 to 0.4 and small *β* (< 0.1). Design I with hyperbolic regulation suggests “fast” effective modification rates of 0.5 to 1.0*hr*^−1^ and a “slow” modification rate less than 1/10 of these values. Presumably, similar to the KaiABC clock, slow conformational dynamics of the protein(s) structure is implicated in the slow regulatory (and period-determining) dynamics. A further prediction of these models is the rather general approximate power-law dependence of the oscillation period on the slow rate, which might be tested using various clock mutants with “cloistered” regulatory site(s). Perhaps these and similar models will assist and encourage the experimental search for additional post-translational protein-based oscillators beyond the remarkable KaiABC cyanobacterial clock.

### A. Methods

Numerical integration of the ODEs was implemented in Fortran (GNU Fortran G77, Free Software Foundation) using 4th-order Runge-Kutta. The numerical routines were also verified in Matlab (The MathWorks, Inc., Natick, Massachusetts, United States); sample scripting code is given in the supplemental section for reproducing oscillations from these models. The sequestration constraint (eqn 5) was implemented using a threshold of 0.001 for 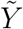. In-house fortran code was written for scanning parameter space, peak finding, estimating oscillatory periods (typical uncertainty < 0.05*hr*), and applying perturbations (fig 8). For figures 7 and 8, the initial conditions, *x*_1_(*t* = 0) = 0.5 = *x*_2_(*t* = 0) were used. Other parameters are listed in the main text. Colormap figures (Fig 3B,D, Figs 6,9) were generated in Matlab (R2016a).

## ACKNOWLEDGMENTS

I appreciate comments and discussion on early work on this project from Carl Johnson, Tetsuya Mori, Ximing Qin, and Yao Xu (Vanderbilt U.). I also express my appreciation to Spring Hill College for supporting faculty research.

## Supplemental Information

### A. Brief derivation of occupancy dynamics in a protein population and selective sequestration of Y

Let *N_X_* be a fixed number of proteins *X* in some volume *V*. Let *N_Y_* be the number of free (non-sequestered) effector molecules in the same volume at time *t*. In the 2-state model let the sites (residues) on each *X* protein be labeled *a* and *b*. Let *N_a_*(*t*) represent the number of residues occupied at site *a* at time *t*. Then *N_X_* − *N_a_*(*t*) is the number of unoccupied *a* sites at time *t*, since the number of X proteins is equal to the number of residues of each type and both are fixed constants. The same notation can be applied to *b*. It is reasonable to assume the rate of change of the occupancy of each site in the population is proportional to the number of unoccupied sites available and depends, by assumption, on the effector molecule (*Y*) in some manner. Allowing for a constant “auto” de-occupancy rate (*k*_−_) from the residue implies a rate of change of occupancy,

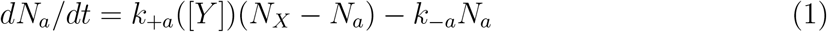

Scaling (dividing) both sides of this equation by *N_X_*, we define the fractional occupancy of site *a* in the population: *x_a_ ≡ N_a_/N_X_*:

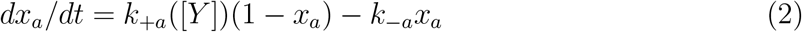

The same analysis applies to site *b*. In the main text Eqn 1, for ease of readability, *a* was replaced by 1 and *b* by 2. This can clearly be generalized to a protein with *k* sites and 2^*k*^ distinct protein states.

Given *N_X_* fixed number of proteins with 2-states, there is no reason to suspect any protein as special so that the distribution of the 4 protein states among the population will follow simple probabilistic arguments and can be computed from the occupancy fractions (at time *t*) given above. Thus if *x_a_* is the fractional occupancy of site *a*, and *x_b_* is the fractional occupancy of site *b*, the probability a given protein has both sites occupied is the product of probabilities of occupancy, *x*_(*a*=1*,b*=1)_ = *x_a_* ∗ *x_b_*; the expected (average) number of proteins at time *t* with both sites occupied is *N_X_* ∗ *x_a_* ∗ *x_b_*. Similarly the expected (average) number of proteins at time *t* with site *a* occupied but site *b* un-occupied is *N_X_* ∗ *x_a_* ∗ (1 − *x_b_*) since 1 − *x_b_* is the probability that site *b* is not occupied. This is the origin of Eqn 2 in the main text. For the purposes of discussion below let *X_k_* with *k* = 1 − 4 represent the 4 protein states such as, *X*_0_ ≡ (0, 0)*, X*_1_ ≡ (1, 0)*, X*_2_ ≡ (0, 1)*, X*_3_ ≡ (1, 1), and *N_k_* represent the number of proteins in each of these states.

It is cumbersome to directly model sequestration of *Y* using mass action, because *Y* must be transiently binding (and un-binding) rapidly from *X* to regulate site occupancy; this is similar (and based on) how KaiA appears to interact with KaiC and affect (auto) phosphorylation in the KaiABC clock ([1], [2]), but KaiA also appears to be sequestered/inactivated by a special state of KaiC, based on KaiB association and distinctive phosporylation states ([3], [4]). The purpose of this paper is to provide a simpler phenomenological model which requires a minimal number of state variables, yet retains the essential oscillatory mechanism(s). Even in the simplest version where one of the 4 protein states of *X* inactivates/sequesters *Y*, brute-force mass action needs to include all possible state transitions among the 4 unbound protein states (*X_j_*), the formation of all possible transiently bound protein complexes with *Y* (the *X_j_* +*Y ↔ X_j_Y* states), the relevant state transitions among these states, allowing for varying rate constants for association/dissociation Y from each of these states and allowing for potential variations in the rate of site modification (e.g., phosphorylation) for each state. Needless to say, the number of differential equations needed for brute-force mass-action to more precisely model the dynamics of the system is much larger than two, resulting in a much enlarged state space and parameter space. This program of mass-action has been carried through in detail for a protein with 4 states (2 sites), and kinase and phosphatase enzyme modifiers in the excellent numerical study of selective sequestration by Jolley et al. [5]; using mass action resulted in 14 ODEs from a the 4-state system and the need for statistical sampling of the large parameter space for finding oscillatory regimes and oscillatory designs/motifs. A simpler assumption that significantly reduces the state space of variables is to assume the rate at which *Y* is sequestered (or released) is directly proportional to the rate at which the sequestering state forms (or degrades):

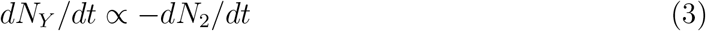

Assuming proportionality of these rates implies rapid sequestration and release of *Y* (much faster than the circadian timescale) based on the instantaneous number of proteins (*N*_2_) in the sequestering state, *X*_2_. The proportionality constant specifies the relative number (*n*) of Y molecules sequestered per protein in the state *X*_2_ = (0, 1) so that by direct integration,

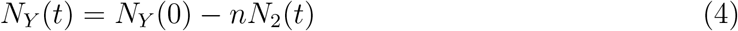

Noting that the number of proteins in the sequestering state is *N*_2_ = *N_X_x*_(0,1)_ and dividing by the presumed fiducial volume (and neglecting molar mass differences) gives Eqn 4 in the main maunscript:

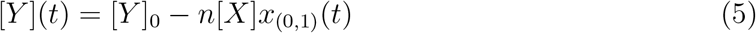

Finally non-dimensionalizing indicates the relevant regulatory parameter is the initial ratio of concentrations, [*Y*]_0_ to [*X*], where there there is no requirement for multiple sequestration of Y by each X (e.g.,one can set *n* = 1) except to lower the effective intial ratio of concentrations:

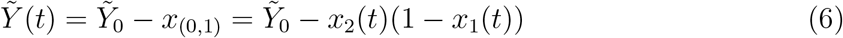

where 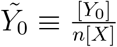. In the main text, 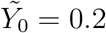 in Fig2A and 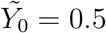 in Fig4A.

### B. Linear Perturbative Analysis of Simple Transferase Model

In this section a linear perturbative analysis about the steady-state solutions of the design II models (Eqn 7) is used to suggest constraints on the parameters that permit (or do not permit) limit cycle oscillations. for convenience the model equations for design II are copied below:

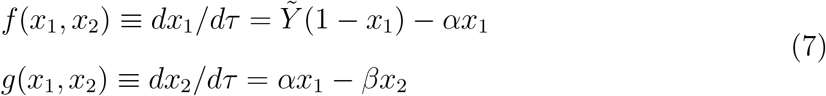

The fixed points 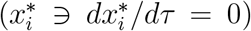 are solvable analytically (from a cubic) and are functions of two effective parameters only, *β* and 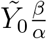. A linear perturbation about the steady-states yields a stability matrix, evaluated at the steady state(s):

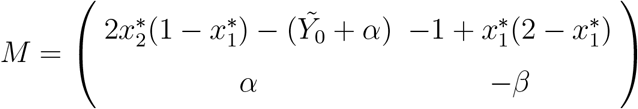

**FIG. 1.**
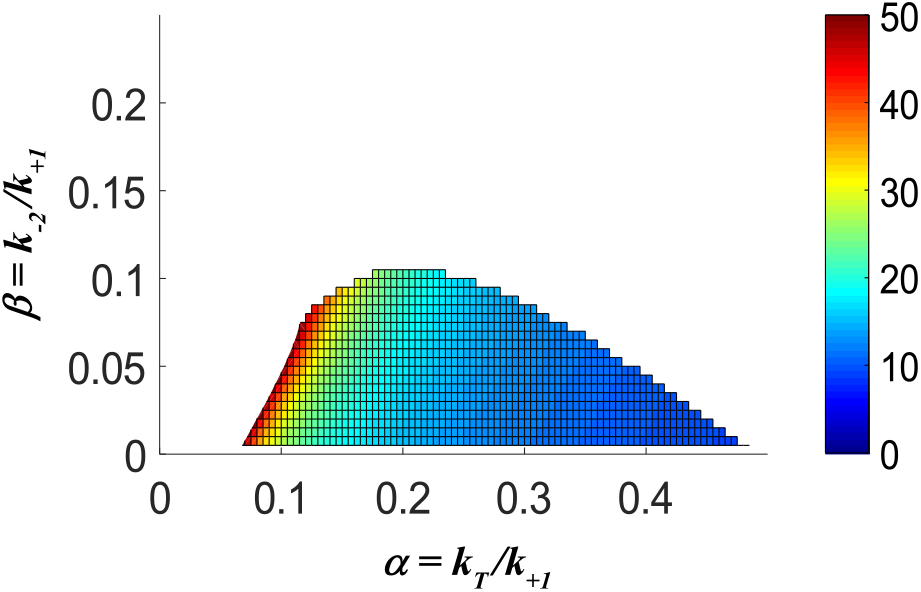
Predicted oscillatory parameter space and dimensionless oscillatory period (colormap) for design II using analytical solutions for the fixed point(s) of the dynamical system (eqn 7) and applying the conditions for (linear) instability of the steady-state(s) from the matrix, *M*.

Potential oscillatory solutions with instability of the steady state (unstable spiral) requires *TrM* > 0 and 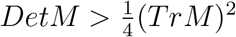. Since 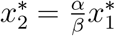 and 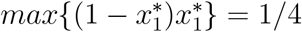 this implies

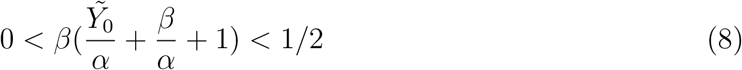

Since each term is positive this implies *β* < 1/2. Further constraints are given using Bendixson’s theorem; the sum of partial derivatives is

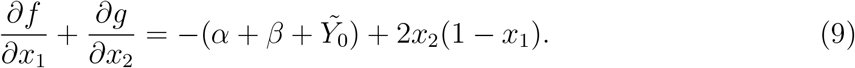

As the latter quantity is confined on the interval [0, 2] assuming the domain 0 < *x_i_* < 1 this sum of partials will be strictly negative (and thus not permit limit cycles) unless

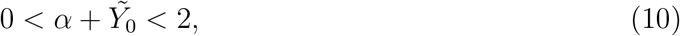

Since each parameter is positive, *α* < 2, 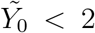. Direct numerical evaluation of the steady states and applying the conditions for instability as functions of *α* and *β* give tighter ranges of instability consistent with these analytical constraints (Supplemental Fig 1). The imaginary component(*ω*) of the eigenvalues of *M* (for each unstable steady-state from solving the cubic and applying the instability criteria) gives a “local” perturbative approximation to the dimensionless period 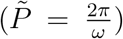 as functions of *α* and *β* (see Supplemental Fig 1 colormap). The estimate of the linear perturbative analysis is in good agreement with the numerical estimate of the oscillatory parameter space and corresponding oscillation periods from direct numerical integration of the ODEs.

### C. Sample Matlab Scripting Code for verifying oscillations

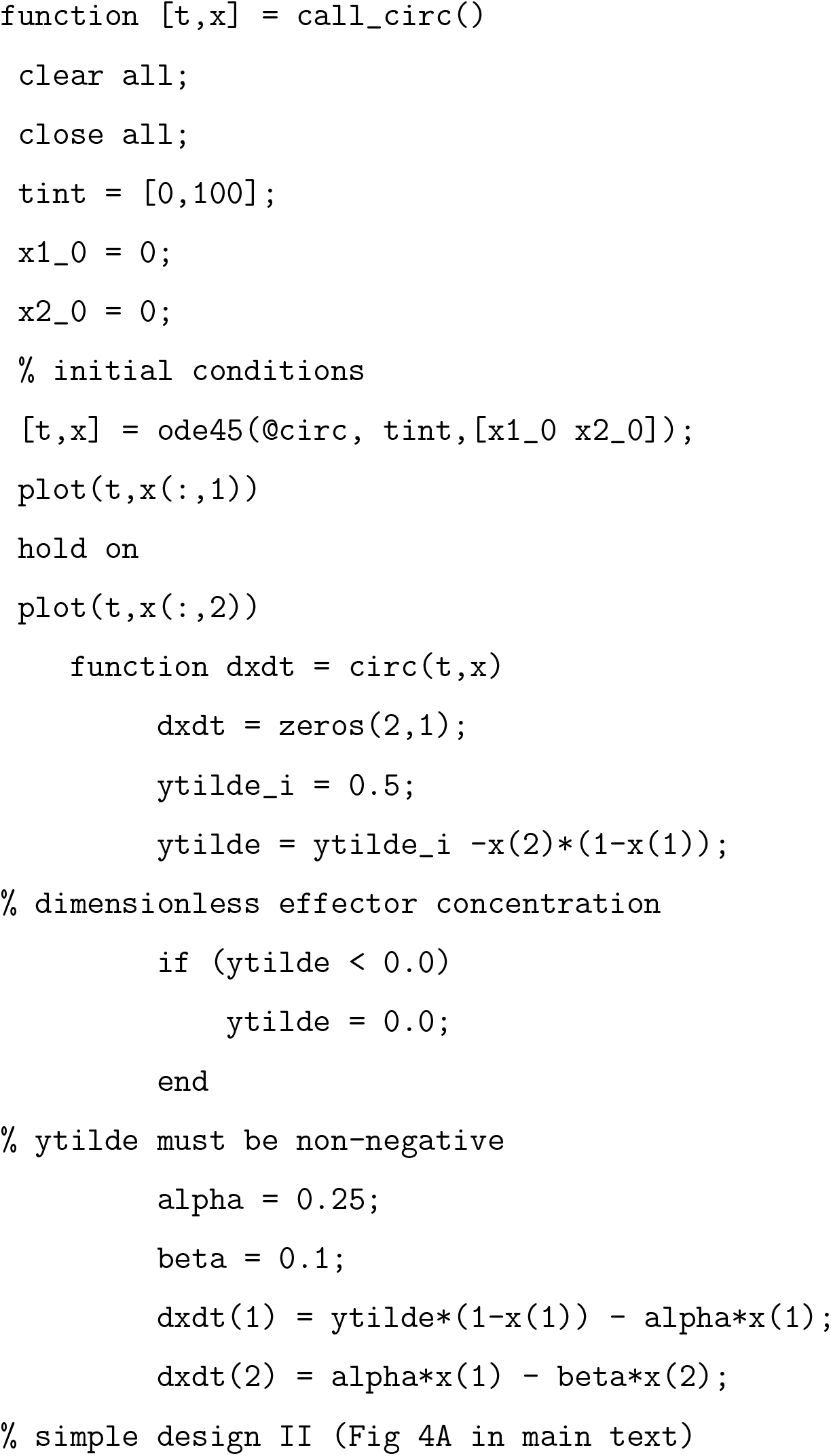
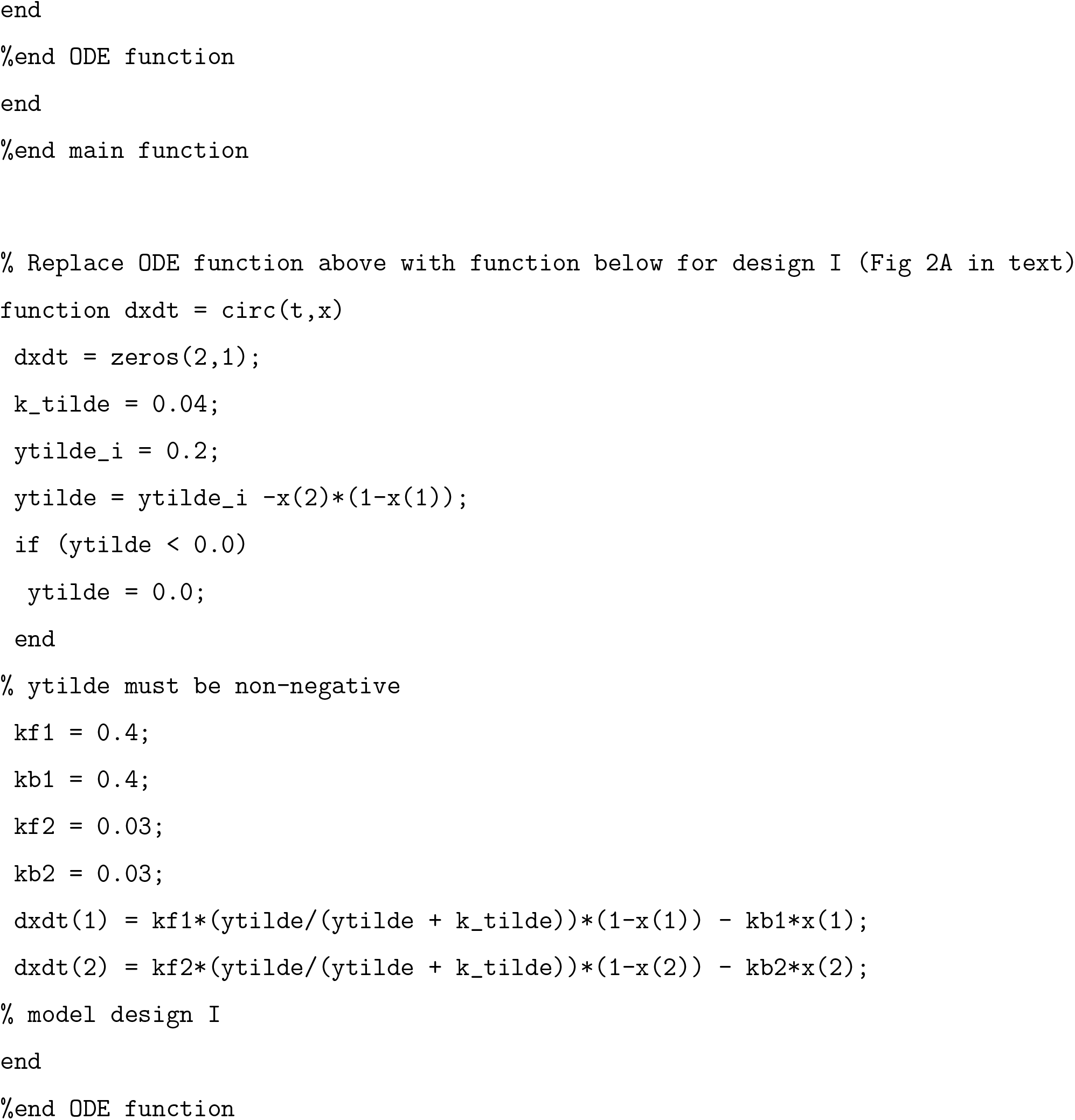

## Notes

### Competing Interest Statement

The authors have declared no competing interest.

### Summary of Updates

Supplemental section included with sample matlab scripting code provided for verifying model oscillations from equations in main text, expanded derivations of equations in main text and additional references. Various typos were also corrected in the main text.

## References

[1] J. C. Dunlap, Cell 96, 271 (1999).

[2] T. Roenneberg, S. Daan, and M. Merrow, Journal of Biological Rhythms 18, 183 (2003).

[3] C. I. Hong, E. D. Conrad, and J. J. Tyson, Proceedings of the National Academy of Sciences 104, 1195 (2007).

[4] J. Tomita, M. Nakajima, T. Kondo, and H. Iwasaki, Science 307, 251 (2005).

[5] M. Nakajima, K. Imai, H. Ito, T. Nishiwaki, Y. Murayama, H. Iwasaki, T. Oyama, and T. Kondo, Science 308, 414 (2005).

[6] M. W. Young and S. A. Kay, Nature Reviews Genetics 2, 702 (2001).

[7] C. Lee, J.-P. Etchegaray, F. R. Cagampang, A. S. Loudon, and S. M. Reppert, Cell 107, 855 (2001).

[8] P. L. Lowrey and J. S. Takahashi, Annual review of genetics 34, 533 (2000).

[9] G. Kurosawa, A. Mochizuki, and Y. Iwasa, Journal of theoretical biology 216, 193 (2002).

[10] X. Qin, M. Byrne, Y. Xu, T. Mori, and C. H. Johnson, PLoS biology 8, e1000394 (2010).

[11] A. Goldbeter, Proceedings of the Royal Society of London. Series B: Biological Sciences 261, 319 (1995).

[12] J. J. Tyson, C. I. Hong, C. D. Thron, and B. Novak, Biophysical journal 77, 2411 (1999).

[13] D. B. Forger and C. S. Peskin, Proceedings of the National Academy of Sciences 100, 14806 (2003).

[14] J. S. O’Neill and A. B. Reddy, Nature 469, 498 (2011).

[15] J. S. O’Neill, G. Van Ooijen, L. E. Dixon, C. Troein, F. Corellou, F.-Y. Bouget, A. B. Reddy, and A. J. Millar, Nature 469, 554 (2011).

[16] R. S. Edgar, E. W. Green, Y. Zhao, G. van Ooijen, M. Olmedo, X. Qin, Y. Xu, M. Pan, U. K. Valekunja, K. A. Feeney, et al., Nature 485, 459 (2012).

[17] C. C. Jolley, K. L. Ode, and H. R. Ueda, Cell reports 2, 938 (2012).

[18] H. Iwasaki, T. Nishiwaki, Y. Kitayama, M. Nakajima, and T. Kondo, Proceedings of the National Academy of Sciences 99, 15788 (2002).

[19] M. J. Rust, J. S. Markson, W. S. Lane, D. S. Fisher, and E. K. O’Shea, Science 318, 809 (2007).

[20] S. Clodong, U. Dühring, L. Kronk, A. Wilde, I. Axmann, H. Herzel, and M. Kollmann, Molecular systems biology 3 (2007).

[21] J. S. van Zon, D. K. Lubensky, P. R. Altena, and P. R. ten Wolde, Proceedings of the National Academy of Sciences 104, 7420 (2007).

[22] P. Ruoff, Journal of Interdisciplinary Cycle Research 23, 92 (1992).

[23] K. Terauchi, Y. Kitayama, T. Nishiwaki, K. Miwa, Y. Murayama, T. Oyama, and T. Kondo, Proceedings of the National Academy of Sciences 104, 16377 (2007).

## Supplemental References

[1] H. Kageyama, T. Nishiwaki, M. Nakajima, H. Iwasaki, T. Oyama, and T. Kondo, Molecular Cell 23, 161 (2006).

[2] T. Mori, S. Sugiyama,, M. Byrne, C. H. Johnson, T. Uchihashi, and T. Ando, Nature Communications 9, 3245 (2018).

[3] M. J. Rust, J. S. Markson, W. S. Lane, D. S. Fisher, and E. K. O’Shea, Science 318, 809 (2007).

[4] C. Brettschneider, R. Rose, S. Hertel, I. Axmann, A. Heck, and M. Kollmann, Mol Syst Biol 6 (2010), 10.1038/msb.2010.44.

[5] C. C. Jolley, K. L. Ode, and H. R. Ueda, Cell reports 2, 938 (2012).

